# Modulations of right hemisphere connectivity in young children relates to the perception of spoken words

**DOI:** 10.1101/2023.03.03.530976

**Authors:** Kartik K. Iyer, Nicola Bell, David A. Copland, Wendy L. Arnott, Wayne J. Wilson, Anthony J. Angwin

## Abstract

The early school years shape a young brain’s capability to comprehend and contextualize words within milliseconds of exposure. Parsing word sounds (*phonological* interpretation) and word recognition (enabling *semantic* interpretation) are integral to this process. Yet little is known about the causal mechanisms of cortical activity during these early developmental stages. In this study, we aimed to explore these causal mechanisms via dynamic causal modelling of event-related potentials (ERPs) acquired from 30 typically developing children (ages 6 to 8 years) as they completed a spoken word-picture matching task. Source reconstruction of high-density electroencephalography (128 channels) was used to ascertain differences in whole-brain cortical activity during semantically “congruent” and “incongruent” conditions. Source activations analyzed during the N400 ERP window identified significant regions-of-interest (*p*_FWE_<.05) localized primarily in the right hemisphere when contrasting congruent and incongruent word-picture stimuli. Dynamic causal models (DCMs) were tested on source activations in the fusiform gyrus (rFusi), inferior parietal lobule (rIPL), inferior temporal gyrus (rITG) and superior frontal gyrus (rSFG). DCM results indicated that a fully connected bidirectional model with self-(inhibiting) connections over rFusi, rIPL and rSFG provided the highest model evidence, based on exceedance probabilities derived from Bayesian statistical inferences. Connectivity parameters of rITG and rSFG regions from the winning DCM were negatively correlated with behavioural measures of receptive vocabulary and phonological memory (*p*_FDR_<.05), such that lower scores on these assessments corresponded with increased connectivity between temporal pole and anterior frontal regions. The findings suggest that children with lower language processing skills required increased recruitment of right hemisphere frontal/temporal areas during task performance.

## Introduction

The study of cognitive-linguistic processing via the measurement of event-related potentials (ERPs), particularly the N400 component, has been the subject of a plethora of electroencephalography (EEG) research, and has shown to be a generally robust phenomenon that is sensitive to various aspects of cognitive and semantic processing (Kutas & Federmeier, 2011; Skeide & Friederici, 2016). The N400 is evident at a young age, with research demonstrating that as early as the third grade (typically between ages 7 to 9), children exhibit adult-like processing at a semantic and phonological level (Coch, 2015). The temporal sensitivity of the N400, therefore, can provide an online (or real-time) view of semantic processing in developing cohorts. Consistent with the notion that the N400 provides an index of lexical-semantic processing, numerous studies have demonstrated correlations between the N400 and behavioural measures such as vocabulary size (Coch & Benoit, 2015; Panda, Emami, Valiante, & Pang, 2020; Stites & Laszlo, 2017) and listening comprehension (Henderson, Baseler, Clarke, Watson, & Snowling, 2011).

Whilst such research has contributed to the understanding of lexical-semantic processing development in children, the interplay and causality of cortical communication between key language regions within the N400 timescale are yet to be elucidated. Here, effective brain connectivity techniques can provide critical insights. These techniques reveal the directionality and strength of cortical communication between brain regions, providing quantitative descriptors for causal mechanisms underpinning the timescale of cortical activity. To date, language-based neuroimaging studies in children which utilise these techniques are limited, though some have used dynamic causal modelling (DCM) with fMRI data to investigate reading skills in children (Cao, Bitan, & Booth, 2008; Liu et al., 2010; Morken, Helland, Hugdahl, & Specht, 2017). Such research has provided critical insights into differences between children with and without dyslexia in the development of effective connectivity, particularly relating to the inferior frontal gyrus and occipito-temporal gyrus (Morken et al., 2017) and deficits in effective connectivity from the fusiform gyrus to the middle temporal gyrus in children with reading disability (Liu et al., 2010).

Despite such focused work, developmental neuroscientists are yet to fully utilize techniques such as DCM in conjunction with EEG/ERP recordings of the N400. Previous fMRI research exploring spoken word-picture matching in a large cohort of children (ages 5 to 18) showed that increased task-related activation was evident within the ventral visual pathway, including the fusiform, inferior temporal and occipital gyri bilaterally, whereas decreased activation associated with task performance was observed within the inferior frontal gyrus bilaterally (Schmithorst, Holland, & Plante, 2007). The application of DCM to measures of the N400 during a similar task could further elucidate on the effective connectivity present in children during the development and processing of language skills. Such research is particularly important, given that the trajectory of development in lexical and auditory (phoneme) processing of words coincides with an orchestration of key cortical regions which parse incoming visual and auditory stimuli to ascribe semantic meaning (Skeide & Friederici, 2016; Yvert, Perrone-Bertolotti, Baciu, & David, 2012).

With these insights in the foreground, the present study investigated effective connectivity of typically developing school-aged children during semantic processing via an event-related dynamic causal modelling approach (DCM-ERP). We explored whether the associations between effective connectivity of ERP activity within the N400 window would be modulated by localized cortical activity within the fusiform gyrus, parietal, inferior temporal, and frontal gyrus regions when matching spoken words to pictures. Using source localization techniques of EEG and the DCM-ERP model, we examined connectivity modulations of cortical activity between significant source-activated regions during task completion. Parameters from the DCM-ERP model were subsequently tested for associations with behavioural measures of receptive vocabulary and phonological memory, to uncover links between cortical activity strengths and developing language skills.

## Methods

### Participants

Thirty typically developing children (15 females, 3 left-handed, mean age = 7.68 years (standard deviation, SD = 0.77)) were recruited as part of a broader study (Bell, Angwin, Arnott, & Wilson, 2019; Bell, Angwin, Wilson, & Arnott, 2019). As reported elsewhere (Bell, Angwin, Arnott, et al., 2019), all participants used spoken English as their native and primary form of communication, and had nonverbal reasoning at or above the 16^th^ percentile as assessed by the Raven’s Colored Progressive Matrices (Cotton et al., 2005; Raven, 2000). As a guide and as per Australian school grading, five participants were in Grade 1 (aged 5 to 6), sixteen were in Grade 2 (aged 6 to 7), and nine were in Grade 3 (aged 7 to 8) at the time of testing. All tests were administered in a sound-treated room and scored by a qualified speech-language pathologist. Ethical approval for the study was obtained from the Behavioural and Social Sciences Ethical Review Committee at the University of Queensland. All parents gave written informed consent for their children to participate in the study.

### Behavioural measures

For this study we included two behavioural measures: receptive vocabulary and phonological memory. A full description of these behavioural assessments has been previously reported (Bell, Angwin, Arnott, et al., 2019; Bell, Angwin, Wilson, et al., 2019). Receptive vocabulary was assessed using the Peabody Picture Vocabulary Test – 4th edition (PPVT-IV; (Dunn & Dunn, 2007). In this task, children are presented with four pictures and are asked to point to the one corresponding to a word spoken by the test administrator. For the purposes of analysis, raw scores were converted to standard scores.

Phonological memory was assessed using the Children’s Test of Nonword Repetition (Gathercole & Baddeley, 1996). In this test, children are asked to repeat a nonword spoken by the test administrator, with test items divided equally into two-, three-, four- and five-syllable items. Nonword repetition has been found to reflect phonological processing skills and is a significant predictor of reading ability in children (Wagner, Torgesen, & Rashotte, 1994). Using normative data supplied by Gathercole et al. (1996), raw scores on this test were converted into standardised z scores.

### Experimental Design and High-density EEG acquisition

As soon as children were comfortably seated and relaxed, we set up and recorded EEG. We acquired high-density EEG data with the Electrical Geodesics EEG system (GES 300) using a 128 channel HydroCel Geodesic Sensor net at a sampling rate of 500 Hz (Electrical Geodesics, Inc, EGI, Eugene, OR, USA). Impedances were kept below 50 kΩ throughout testing, which is considered acceptable when using high-impedance amplifiers (Ferree, Luu, Russell, & Tucker, 2001). Event-related potentials (ERPs) were generated during semantically congruent and incongruent word-picture task conditions. Children were positioned approximately 0.5 to 1 metre in front of two speakers in a sound-treated and radio-frequency-shielded audiometric booth. Participants were instructed that they would hear a spoken word (via the speakers), and that this auditory stimulus would be followed by a picture in the centre of their computer screen. Children were asked to judge whether the picture was a correct match for the spoken word or not, and to respond verbally with “yes” or “no” when a response prompt appeared on the screen (after the picture presentation). Verbal responses were not made until greater than 1000ms had elapsed to ensure that motor planning related responses were minimal. Auditory stimuli were delivered using e-prime 2.0 presentation software (E-Prime 2.0), with stimuli output levels peaking in the range of 54 to 56 dB for this cohort. A short practice task was presented prior to the actual experiment to ensure children understood the nature of the task and to provide them with practice at waiting for the response prompt. Performance was subsequently monitored throughout the experiment proper to ensure children complied with the instructions. Each trial followed a standardized sequence in timing and presentation of stimuli (**Fig. 1A)**, wherein: (i) a centred fixation cross was displayed on screen for 500 ms (ii) a 2000ms blank screen during which a spoken word was presented via speakers (all spoken word stimuli designed to offset at 1000ms), (iii) a color picture centred on screen displayed for 1000 ms, (iv) a blank screen for 500 ms, to further account for response delays and (v) a question mark centred on screen to prompt the child to make a verbal “yes”/”no” response.

**Figure 1.**
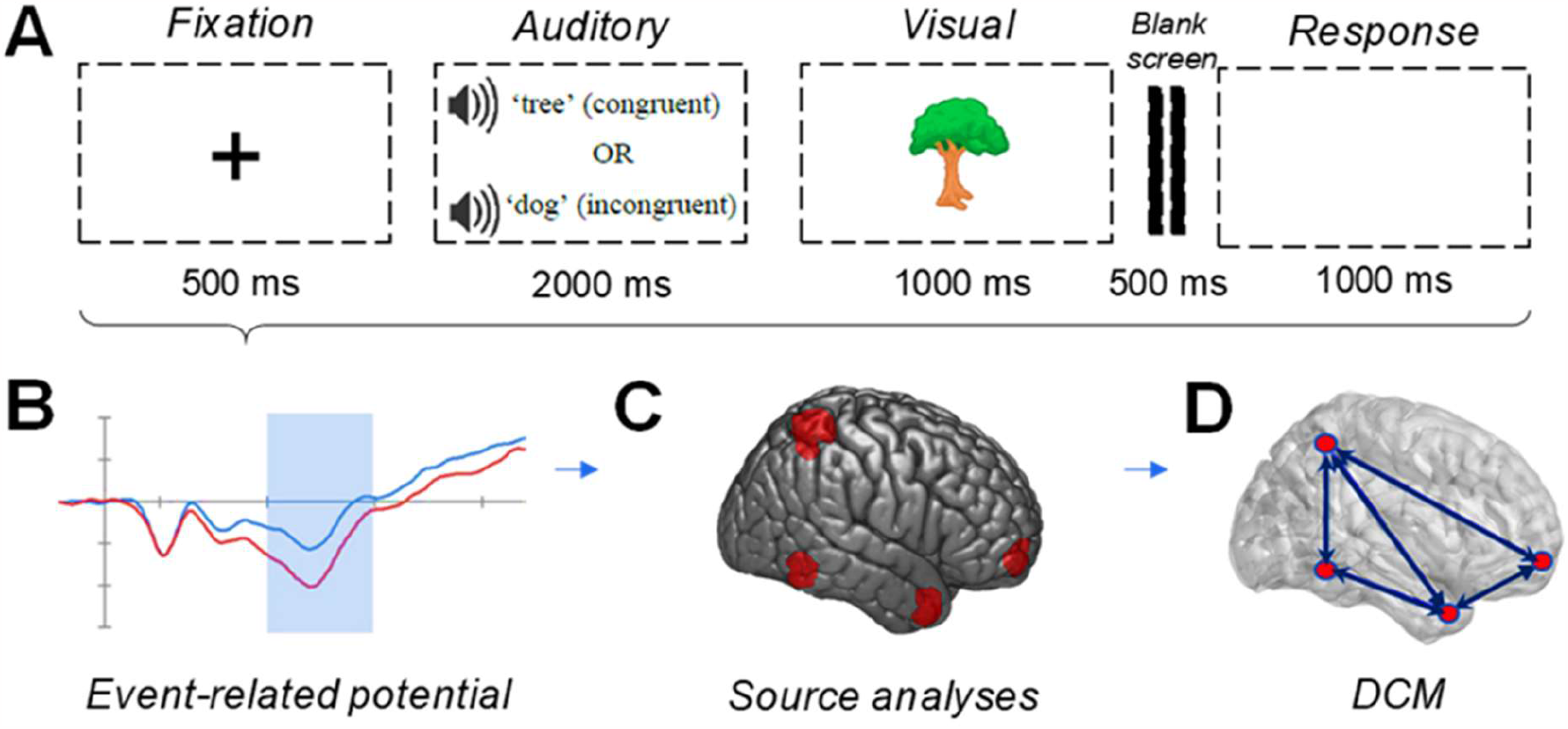
Paradigm and Processing Pipeline. (A) Task paradigm (B) Event-related potentials (ERPs) for each condition (see Bell et al., 2019) (C) Source localization and statistical parametric mapping (D) Dynamic Causal Modelling (DCM) and model hypothesis testing

### Source imaging of EEG

Data was first pre-processed using Netstation EEG software to identify and remove artefacts. EEG segments contaminated by muscular artefacts (e.g., high frequency noise signatures greater than 20Hz) were marked for removal. Further, a visual inspection procedure was then employed to identify and remove any additional EEG artefacts, such as high amplitude data (>200uV) and vertical or horizontal inflections in electrooculogram amplitude that differed by greater than 140uV (Bell, Angwin, Arnott, et al., 2019). Pre-processed EEG data was then processed offline using the SPM12 toolbox (Litvak et al., 2011) and custom scripts in MATLAB (Mathworks, Natick, MA). All data epochs were band-pass filtered between 0.1 and 30 Hz, down sampled to 250 Hz and epoched between -100 ms and 800 ms (centred around stimulus onset at zero seconds). Eye-blinks (measured via electrooculogram channels) and muscular artefacts were removed from ERP trials and bad channels were replaced with data interpolated from surrounding channels. Following these procedures, two participants were subsequently excluded from analysis due to excessive artefacts which resulted in <20 trials available for analysis per condition for those participants. Following EEG data cleaning procedures, we had on average, 32% of congruent trials and 35% of incongruent trials removed per subject, with at least over half of all experiment trials completed per subject available for analysis. All channels were then average referenced and baseline correction was performed. Only ERPs with correct responses to stimuli were included in our analyses. Following removal of incorrect trials and artefacts, an average of 38 (out of 60) congruent trials, and 37 (out of 60) incongruent trials were available for analysis (**Fig. 1B**). All ERPs were then grand averaged for each participant within each condition. Full ERP results based on the scalp-EEG activity are summarized in our recent study (Bell, Angwin, Arnott, et al., 2019).

Source reconstruction of all ERP data on the remaining 28 participants was next performed with in-built 3D source reconstruction routines within SPM12. Given the age bracket of our participants it was particularly important to derive source reconstructed estimates from age-specific MRI priors. Here, we used asymmetric MRI templates between 4.5 and 18.5 years of age from the NIH MRI study of Normal Brain Development (Evans & Group, 2006; V. Fonov et al., 2011; V. S. Fonov, Evans, McKinstry, Almli, & Collins, 2009). Next, we generated an age-specific “pediatric” structural head-model with this dataset which allowed us to account for variations in head sizes and volume differences present in 6-to 8-year-old children. For example, children aged 6 years of age were modelled on MRI template priors in this age bracket alone, with the same approach for participants aged 7 and 8. Using a forward model approach, electrode positions were co-registered with this pediatric head-model template, using a “fine” mesh composed of 20484 vertices and projected into MNI stereotactic space. The forward model consists of a boundary element model (BEM) composed on three layers (scalp, outer skull, inner skull), using a multiple sparse priors approach (Friston et al., 2008). Source imaging was performed on EEG activity following presentation of the visual picture. Cortical source images of ERPs were then generated within the N400 window (300 – 500 ms) to probe specific semantic processes related to this task (Lau, Phillips, & Poeppel, 2008). Early ERP components (150 – 250 ms) were also modelled via source imaging to ascertain cortical areas engaged during early exposure to semantic tasks within the paradigm studied.

### Dynamic causal modelling (DCM)

We used a DCM for ERP approach within SPM12 to measure effective connectivity between cortical sources during the N400 time window. Briefly, the DCM approach employs a neurobiologically generated neural mass model which consists of distributed, laminar neural architectures to denote excitatory pyramidal cells, spiny stellate and inhibitory interneurons. This canonical neural architecture provides a basis for hypothesis testing of underlying changes in direction and strength of cortical activity, as induced by experimental conditions, between different source regions. Specifically, for the DCM approach we used the ERP model architecture, which is a modified version of the Jansen-Rit (David et al., 2006) which enables us to measure modulations present within ERP waveforms elicited during task-states. Our model space for DCM was guided by selection of specific regions-of-interest (ROIs) identified by statistical parametric mapping results from our source localization analyses (see *Results*, **Fig. 2**). Using these ROIs, we constructed a model space testing 48 different hypotheses categorized by the following model families: (1) forward connections only (2) backward connections only and (3) forward and backward connections (fully connected, bidirectional). Further model variations within each of these model families included the absence or presence of self-connections (intrinsic connections) on single ROIs (e.g. self-connection on rFusi); and all possible presence of multiple self-connections (i.e. self-connections modelled on rFusi, rIPL and rSFG simultaneously). Each of these 48 models underwent model inversion via DCM. Our model inversion window was fixed to 0 to 600 ms to include early ERP components with purported N400 components, to model early and late language cortical processing.

**Figure 2.**
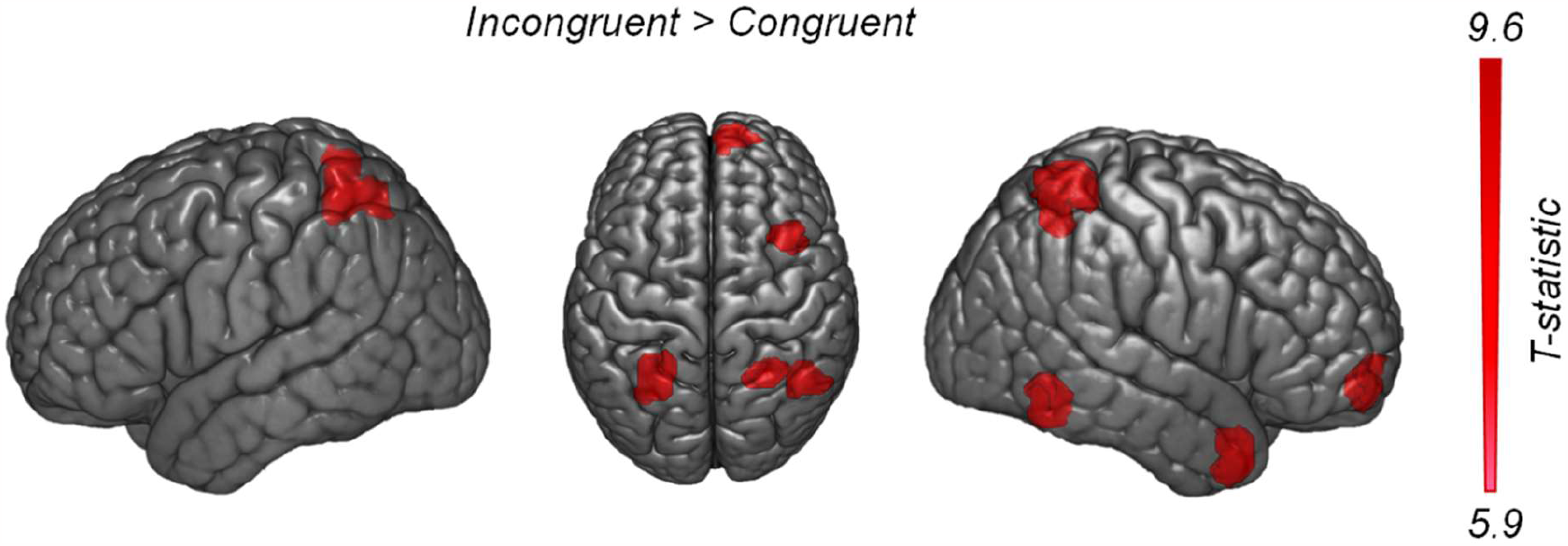
Source localization of event-related potential in N400 window. Contrasts of semantically-incongruent versus semantically-congruent pairs indicated four key ROIs activated in the right hemisphere: (rFusi, rIPL, rITG, rSFG), and one left hemisphere ROI (lIPL). Sagittal and axial views shown to capture all ROIs that survived significance testing.

### Statistical analysis

#### Statistical parametric maps

Group level contrasts of N400 ERP source images were examined within SPM12 using a random-effects analysis. Here, one-sample t-tests were first examined to assess source activations within semantically congruent and semantically incongruent conditions, respectively. We next tested the hypothesis that source activations of the early (150 to 250 ms) window and the N400 within the semantically incongruent condition would be greater than the congruent condition, and, vice versa, that congruent would be greater than incongruent. Results from these statistical contrasts (statistical parametric maps, **Fig. 1C)** informed our ROI selection for DCM analysis (**Fig. 1D**). Source activations were identified via cluster-level search thresholds (activations with 10 mm radius, *p* < .001 uncorrected height threshold, and a cluster-level threshold (kE) of 200 voxels). Family-wise error correction (with statistical significance set at *p*_FWE_ < .05) for multiple comparisons was then applied to identify statistically significant source activated regions.

#### Bayesian approach

A random-effects Bayesian model selection (BMS-RFX) was used to derive the exceedance probability of all models and determine the “winning” model (i.e. the model with the highest exceedance probability). The BMS-RFX approach assesses the validity of model generated data for a randomly chosen subject, determining the likelihood of that model for a representative group. Using this approach we also derived exceedance probabilities to test model variants within our DCM framework. This included testing of (i) model families that consisted of fully connected, forward and backward connections and (ii) models with specific self-connections versus those that had no self-connections. Valid DCM connections from the winning model and their coupling strengths, represented in units of hertz (Hz), from each participant were extracted via the exponent of DCM B-matrices. These matrices contain coupling values which provide a measure of the strength of directionality between regions. Here, values above 1 Hz are indicative of increases in excitation between ROIs during tasks, whereas coupling values below than 1 Hz indicate decreases in excitatory influence during task stimulus.

#### General linear regressions with behavioural measures

General linear regression analyses of the association between coupling strengths (dependant variable) and receptive vocabulary (PPVT-IV) and phonological memory (Children’s Test of Nonword Repetition), were performed to test the putative link between cortical connectivity strengths in the winning model with language assessments. Multiple comparisons of all within-group correlations performed between connectivity coupling strengths from all cortical connections within the DCM-B matrix and behavioural measures were adjusted for false discovery rate via the Benjamini-Hochberg test, *p*_FDR_ < .05. Post-hoc tests of DCM connectivity values with age and gender were also checked via general linear regression and a Mann-Whitney rank sum test, respectively.

## Results

### Source localization of event-related potentials

Results from our source reconstruction analyses of event-related potentials in N400 windows (300 to 500 ms) indicated significant clusters activated by task-effects in five key ROIs, namely: the right fusiform gyrus (rFusi), left and right inferior parietal lobule (lIPL and rIPL), right inferior temporal gyrus (rITG) and right superior frontal gyrus (rSFG). Source-based contrasts of early ERP windows (150 to 250 ms) confirmed engagement of all regions aforementioned (**Supplementary Fig. 1A)** except for rSFG. Here, the semantically incongruent condition had a significantly greater task-effect compared with the semantically congruent condition (*t*-test, family-wise error corrected: *p*_FWE_ < .05 with a cluster-level threshold (kE) ≥ 220 and *t*-statistic range: 5.9 to 9.6 for all ROIs, **Fig. 2**). ROIs from **Table 1** indicate that active regions within the N400 window were predominantly featured on the right hemisphere (**Fig. 2**). Comparisons of source activity at the ROI-level of lIPL and rIPL regions from early and later ERP windows indicated similar source estimates observed across all participants (**Supplementary Fig. 1B**).

**Table 1.** Significant source activations during the word-picture matching task. BA= Brodmann’s Area, Montreal Neurological Institute (MNI) coordinates in *x, y, z* space. Regions included: fusiform gyrus (rFusi), inferior parietal lobule (rIPL and lIPL), inferior temporal gyrus (rITG) and superior frontal gyrus (rSFG). *T*-statistics for each region indicated height thresholds, following multiple comparisons correction (*p*_FWE_ < .05, cluster threshold *k*_E_> 300)

| Significant clusters ($p_{\text{FWE}} < 0.05$) | Abbreviation | BA | MNI coordinates | | | $T$-statistic |
| --- | --- | --- | --- | --- | --- | --- |
| | | | $x$ | $y$ | $z$ | |
| Fusiform gyrus | rFusi | 37 | 46 | -56 | -8 | 6.08 |
| Inferior parietal lobule, *right* | rIPL | 7 | 28 | -54 | 62 | 8.77 |
| *left* | lIPL | 7 | -28 | -62 | 60 | 9.60 |
| Inferior temporal gyrus | rITG | 38 | 34 | 12 | -38 | 6.97 |
| Superior frontal gyrus | rSFG | 8 | 6 | 62 | -12 | 5.90 |

**Table 1** – Significant source activations during the word-picture matching task.
| Significant clusters ($p_{FWE} < 0.05$) | Abbreviation | BA | MNI<br>coordinates | | | $T$-statistic |
| --- | --- | --- | --- | --- | --- | --- |
|  |  |  | x | y | z |  |
| Fusiform gyrus | rFusi | 37 | 46 | -56 | -8 | 6.08 |
| Inferior parietal lobule, *right* | rIPL | 7 | 28 | -54 | 62 | 8.77 |
| *left* | lIPL | 7 | -28 | -62 | 60 | 9.60 |
| Inferior temporal gyrus | rITG | 38 | 34 | 12 | -38 | 6.97 |
| Superior frontal gyrus | rSFG | 8 | 6 | 62 | -12 | 5.90 |

### DCM and Bayesian model selection

Inverted DCMs for each individual model were iteratively tested via our model space (**Fig. 3**). The model space was optimized by testing numerous combinations of forward, backward, bidirectional and self-connections modelled on balance with the size of the sample available and rationale for various hypothesis tests (28 participants; Figure 3 contains 15 primary DCM models tested, with Supplementary Fig. 2 consisting of the remaining 33 models tested).

**Figure 3.**
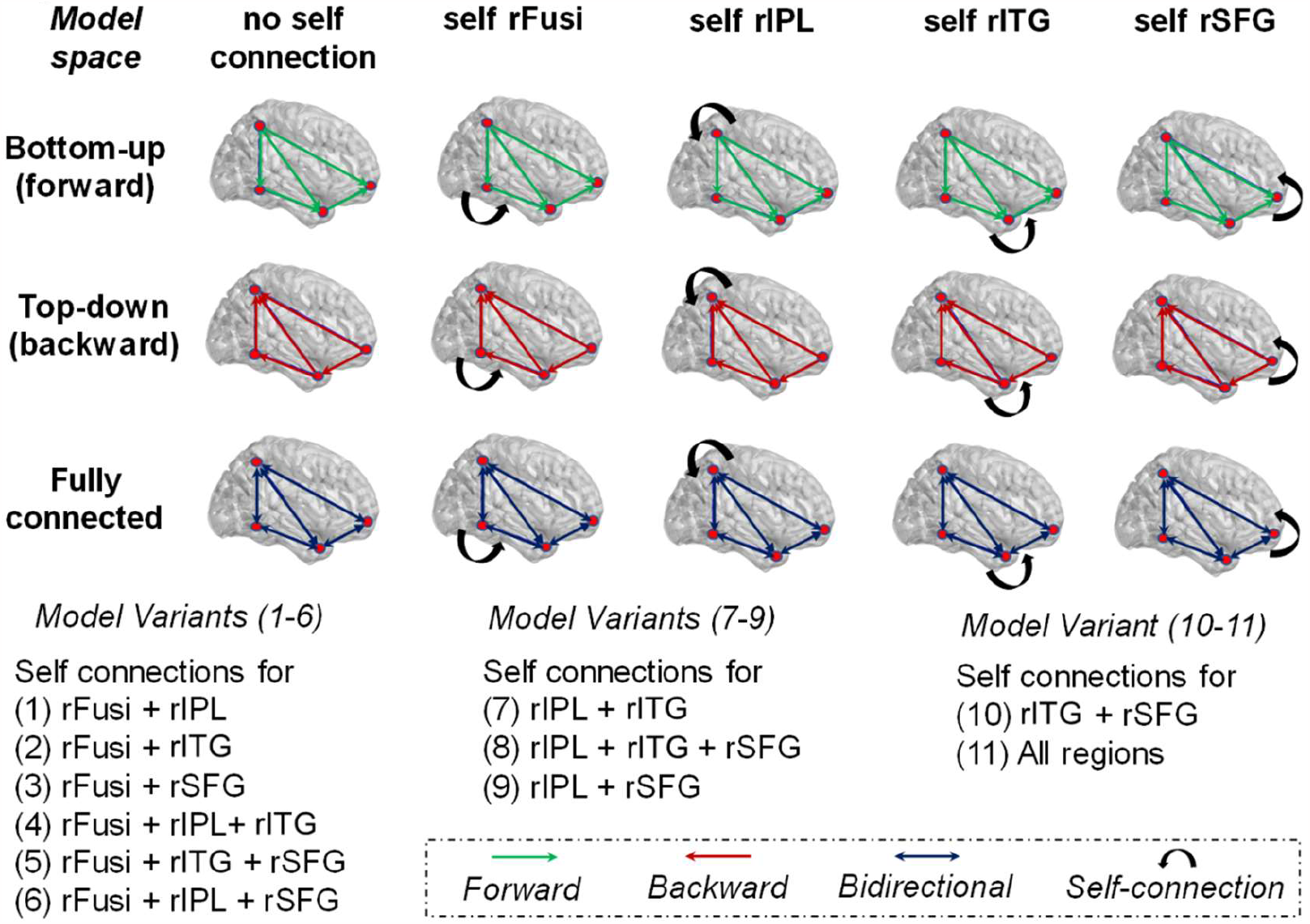
Model space for hypothesis testing via DCM. Here, the basic model space is visualized by three model types: (1) “Bottom-up” (or forward only connections); (2) “Top-down” (or backward only connections); (3) Fully connected (bidirectional connections). Variations of these models are enumerated by the absence/presence of self-connections over various ROIs.

Second-level analyses via BMS-RFX indicated that a fully connected model most likely explained effective connectivity that exists within this group of children for this task (**Fig. 4A**). Here, the model exceedance probability indicated with a high likelihood (**Fig. 4B**) that these models have self-connections on rFusi, rIPL and rSFG nodes. Family-level testing of exceedance probabilities further indicated that within this model scheme, a fully connected bidirectional model was the dominant model type (**Fig. 4C**), and that there was a high likelihood of models consisting of self-connections over the rSFG node, in comparison to models without this self-connection (**Fig. 4D**).

**Figure 4.**
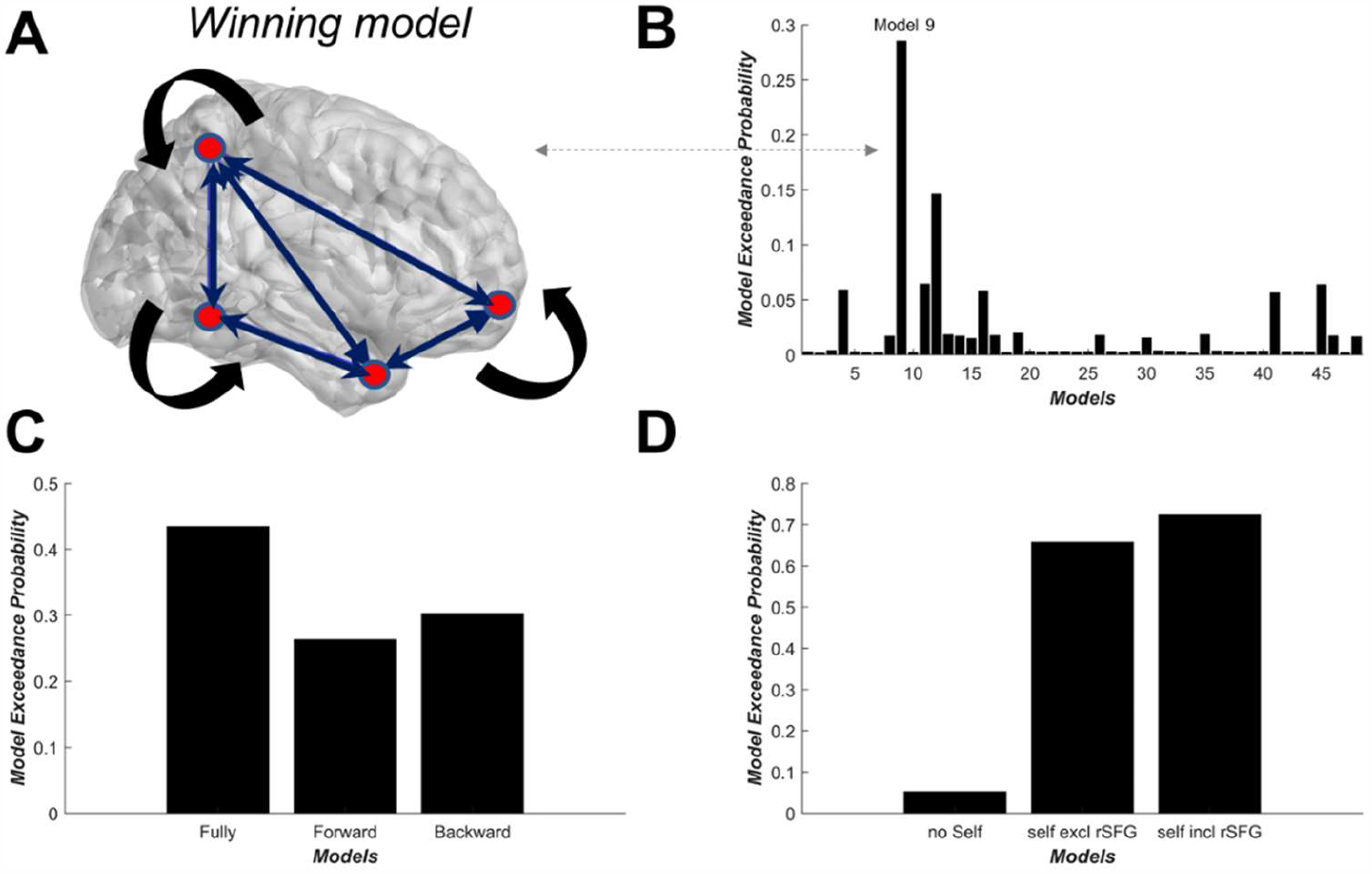
Bayesian model selection and family model effects. (A) The “winning” model as determined by BMS-RFX. (B) Model exceedance probabilities across the 48 models tested, indicated here via the dashed line. Family-level testing via BMS-RFX of (C) model type and (D) the absence versus the presence of self-connections on rSFG.

Correlations of DCM modulatory coupling parameters (DCM-B matrix) from the winning model with behavioural measures of receptive vocabulary and phonological memory revealed significant associations for specific connections. Namely, connections between rITG and rSFG were found to covary with semantic and phonological variables (**Fig. 5**), following adjustment for false discovery rate (Benjamini-Hochberg test). Specifically, decreasing coupling strengths of forward and backward connections from rITG to rSFG were significantly correlated with increasing receptive vocabulary scores (*R*^*2*^ *=* .5, *p*_FDR_ = 0.0003, **Fig. 5A**). Moreover, decreasing coupling strengths of forward and backward connections between rITG and rSFG were both significantly correlated with increasing phonological memory performance (*R*^*2*^ *=* .38, *p*_FDR_ = 0.0063). Primarily, these findings suggest that decreased cortical drive in connections between rITG and rSFG during the task (coupling values < 1) were associated with higher receptive vocabulary and phonological memory scores, contrasted with those children with lower receptive vocabulary and phonological memory, who tended to display an increased cortical drive during the task in these right hemisphere regions (coupling values > 1). Additional tests revealed that connectivity values and age were not significantly associated (R^2^ = 0.05, *p =* 0.27) nor suggested any gender differences (*p* = 0.55 via Mann-Whitney test).

**Figure 5.**
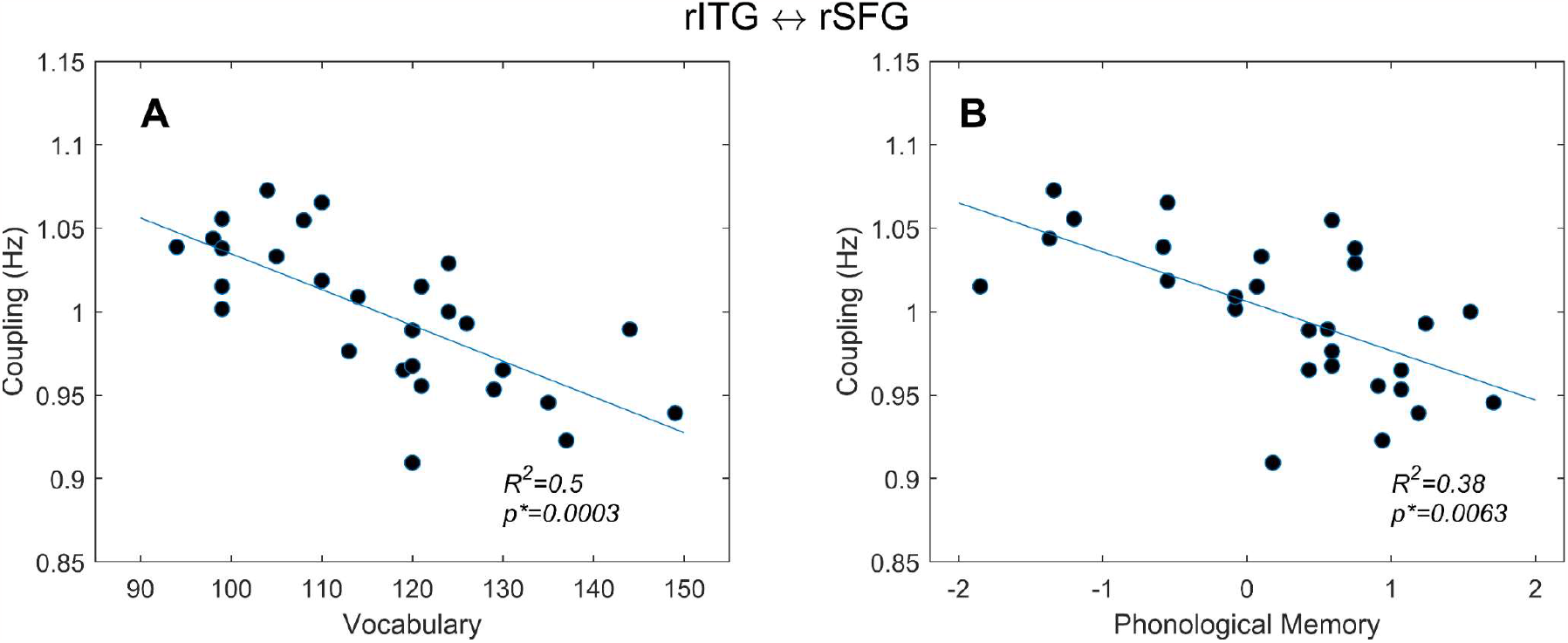
Associations between coupling strengths from winning DCM model with semantic and phonological measures. Increasing scores in these measures corresponded with decreasing coupling strengths between these regions for (A) – (B) forward and backward connection from rITG to rSFG, with each respectively negatively correlated with receptive vocabulary and phonological memory (z-scored). Correlations as measured by general linear modelling (*R*^2^) and tested for significance (*p*_FDR_<.05, asterisked)

## Discussion

The present study examined modulations in cortical effective connectivity in young school-aged children (6 to 8 years of age), during participation in a spoken word-picture matching task with high-density EEG. As revealed by source imaging and DCM, our main findings suggest that changes in cortical effective connectivity during semantic processing were predominantly localized within the right hemisphere. Firstly, source localization effects within the N400 window indicated a coherent network comprising rFusi, rIPL, rITG and rSFG regions, with only one activated region in the left hemisphere (lIPL). In interrogating a DCM model space that focused on the interactions within these right hemisphere ROIs, the winning model indicated a fully bidirectional network, with self-connections over rFusi, rIPL and rSFG areas. The association between specific connections within this winning model, namely modulatory coupling parameters from rITG and rSFG, were found to be negatively correlated with receptive vocabulary and phonological memory measures. Findings from this study indicate that the decreases in cortical coupling within temporal pole and anterior frontal regions corresponded with increasing receptive vocabulary and phonological memory performance, suggesting that proficiency in semantic processing of young school-aged children is indexed by the cortical activity in these key right hemisphere regions.

The predominantly right hemisphere N400 effect observed here is not totally unexpected, given that the N400 is usually more prominent in the right hemisphere in adults (Federmeier & Kutas, 1999; Kutas & Federmeier, 2000) and young children have shown less left-lateralized language comprehension relative to adults (Brauer & Friederici, 2007; Brauer, Neumann, & Friederici, 2008). These neural responses within the right hemisphere are likely to be attributed to robust task-related effects stemming from spoken word-picture paradigms in children. For example, Schmithorst et al. (2007) fMRI study of spoken word-picture matching in children aged between 5 and 18 years showed activations within the right inferior temporal cortex and the right fusiform regions, consistent with our findings. A meta-analysis of fMRI studies of language in children also observed consistent age-related increases in right fusiform activity during semantic processing (Weiss-Croft & Baldeweg, 2015).

Our use of auditory stimuli is also likely to be a contributing factor in observing these right hemisphere activations. In this regard, evidence from MEG studies of the N400 in adults have indicated tonally active bilateral frontal and temporal regions, including inferior and medial frontal gyri and the anterior temporal lobe during semantic judgements to auditory words (Marinkovic et al., 2003) and auditory sentence processing (Maess, Herrmann, Hahne, Nakamura, & Friederici, 2006). Accordingly, the detection of predominantly right hemisphere cortical N400 activation during spoken word-picture matching, whilst surprising, is not without context. The role of these right hemisphere regions may be further explained by our DCM and behavioural results.

Notably, the most prevalent feature of the winning DCM model was the presence of self-connections over fusiform, inferior parietal and superior frontal regions in a fully connected right hemisphere network. The addition of these *intrinsic* connections in a DCM-ERP framework have been postulated to augment interpretation of recurrent cortical activity and connectivity modulations within an evoked response network (Kiebel, Garrido, & Friston, 2007). In our work, recurrent cortical activity in a young developing brain could play an important role in regulating ERP activity within these key cortical regions. For example, N400 cortical activity in the fusiform is likely related to its critical role in processing visual word input (Schmithorst et al., 2007; Visser, Jefferies, Embleton, & Lambon Ralph, 2012) and early components of semantic processing (Schmithorst et al., 2007; Shimotake et al., 2015). However, as word-level processing purportedly traverses the canonical dorsal and ventral routes (Lau et al., 2008), the recurrent cortical activity response within the IPL could relate to attentional processes associated with word learning and semantic representation (Skeide & Friederici, 2016). Within ventral streams of semantic processing, as cortical activity passes through inferior temporal gyri before reaching medial regions of the frontal lobe, such as the SFG, recurrent evoked processes related to receptive vocabulary may speak to the role of higher-order aspects of cognitive control including working memory processes (Sabb, Bilder, Chou, & Bookheimer, 2007) and conflict processing (Hu, Ide, Zhang, & Chiang-shan, 2016; Ye & Zhou, 2009). Indeed, our evaluation of family-based statistical inferences on our DCM model space suggests that the dominant mode of intrinsic activity over the SFG (**Fig. 4D**) could be related to an increased load on executive control networks, reflecting the time required for a young participant to wait for the response prompt during task completion.

In exploring the relationship between coupling parameters from the winning DCM model with indices of receptive vocabulary and phonological memory, interactions between rITG and rSFG regions (**Fig. 5**) further elucidate the causal mechanisms of evoked responses within the right hemisphere that mediate language performance. In the first instance, better receptive vocabulary performance corresponded with decreased forward coupling from ITG to SFG. Interestingly, evidence for such relationships, such as that from Blumenfeld, Booth, and Burman (2006) study of a semantic judgement task with auditory and visual word triplets, suggests that higher or lower accuracy is determined by activity in these key brain areas. For example, accuracy for the auditory version was associated with activation in the right MTG, while lower accuracy was associated with increased frontal activity (IFG/MFG). Similarly, for the visual condition, higher accuracy was associated with activity in the left MTG and right ITG and MFG, and lower accuracy associated with activity in the IFG bilaterally.

For phonological memory, our observation of decreased coupling of forward and backward connections of ITG and SFG was associated with better nonword repetition performance. This is in line with the idea that temporal and frontal lobes are responsible for maintaining components of phonological working memory load (Scott & Perrachione, 2019), where task-related deactivations to nonword discrimination conditions have been specifically observed in bilateral superior temporal and frontal gyri (Perrachione, Ghosh, Ostrovskaya, Gabrieli, & Kovelman, 2017; Scott & Perrachione, 2019). From a broader developmental perspective, these links to receptive vocabulary and phonological memory may also support the idea that children with thicker cortices rely less on right hemisphere language areas to process verbal linguistic information as syntactic processing shifts towards left hemisphere structures (Nunez et al., 2011).

There are several explanations as to why children with lower receptive vocabulary or phonological memory may have increased frontal recruitment in right hemisphere areas, which could in part be attributed to inadequate or inefficient access to semantic and phonological word-level representations. Blumenfeld et al. (2006) suggested that children with worse semantic judgement may have a more weakly connected semantic system, and that anterior areas are more likely recruited for semantic access when there is reduced efficiency of posterior regions linked with lexical-semantic processing. Of interest, Panda et al. (2020) found that a frontal N400 was more prominent in children with weaker receptive vocabulary, whereas children with stronger receptive vocabulary displayed a tendency towards a stronger N400 at central/parietal regions. The authors postulated that the lexico-semantic processes indexed by the N400 may become more efficient and localized with age and the development of receptive vocabulary.

Another potential explanation stems from structural diffusion tensor imaging, where the integrity of tracts within the right hemisphere, such as the arcuate fasciculus and longitudinal fasciculus connecting temporal and frontal regions, relates to word recognition and comprehension (Horowitz-Kraus, Wang, Plante, & Holland, 2014). In consideration of these findings, the results for the present study suggest that children with lower receptive vocabulary and phonological memory scores may have less efficient or less elaborate semantic representations compared to those with more efficient language processing skills. Typically developing children with lower language processing capabilities may need to recruit additional resources in the temporal and frontal areas of the right hemisphere during task performance (Jasińska et al., 2021), resulting in increased cortical drive within these regions.

A limitation of the current study is the absence of strong cortical activations in the left hemisphere, which constrained our overall DCM model space. We opted for our hypothesis driven DCMs to be informed by regions-of-interest identified by our source localization analyses, where strong cortical activations were primarily localized to the right hemisphere. This approach however is two-fold in potential advantages (i) a constrained DCM model space focusing primarily on right hemisphere regions offers a more in-depth interrogation of cortical connectivity across high and low language performers (ii) overall model complexity is optimised. A comparative analysis of the right inferior parietal lobule with its left hemisphere homologue was performed to provide further support that cortical activity within the one significant left hemisphere region (lIPL) operates within similar dynamic ranges to rIPL across ERP timescales (Supplementary Figure 1B). Future approaches could consider expanding the cortical ROIs tested, to interrogate fully bilateral cortical connections (i.e., left and right hemisphere homologues), though a robust source-imaging based rationale is recommended for EEG data in the absence of above-threshold cortical activity in left hemisphere regions.

### Conclusions

In this study, we used DCMs of ERPs to examine language processing in children participating in a spoken word-picture paradigm. We found that cortical N400 responses during the task were localized primarily in the right hemisphere, with specific connectivity modulations from temporal and frontal regions. Increased cortical drive in these regions negatively correlated with receptive vocabulary and phonological memory. Our findings point to the existence of relatively simple causal mechanisms of cortical N400 activity between inferior temporal and superior frontal gyri, thereby highlighting the role of the right hemisphere in modulating cortical activity during key language processes (i.e. semantic access and perception of spoken words) in children.

### Data availability statement

Data are available upon reasonable request addressed to the corresponding author.

## Acknowledgements

The authors thank the families and child participants for their availability and participation in the experiment, and Hear and Say, Brisbane for assistance with recruitment. A. J. A. acknowledges support from the ARC Centre of Excellence for the Dynamics of Language (Project ID: CE140100041). D. A. C was supported by a University of Queensland Vice-Chancellor’s Fellowship.

## Supplementary figures

**Supplementary Figure 1.**
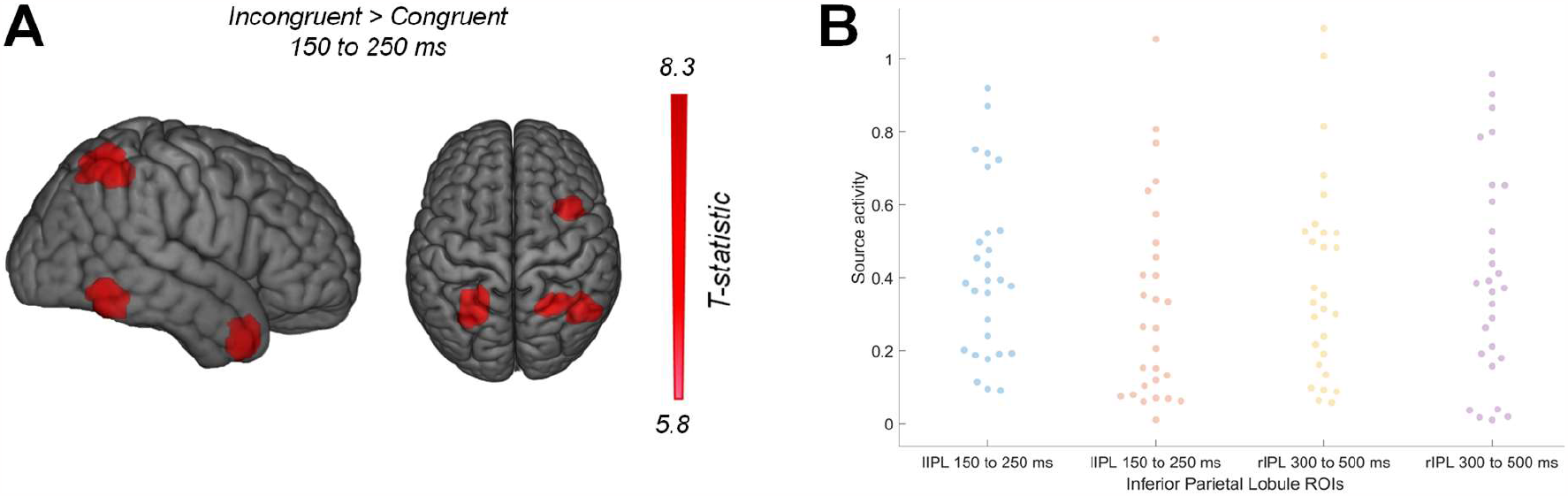
Source localization effects across the ERP timescale. (A) Contrasts of semantically-incongruent versus semantically-congruent pairs during the 150 ms to 250 ms window indicated three key ROIs activated in the right hemisphere: (rFusi, rIPL, rITG), and one left hemisphere ROI (lIPL). Sagittal and axial views shown to capture all ROIs that survived significance testing. (B) A comparison of lIPL and rIPL source activity components at the region-of-interest level indicates that source estimates in both of these regions are highly similar. No significant differences were observed across multiple comparisons of lIPL versus rIPL at both early and later ERP windows (FDR testing applied).

**Supplementary Figure 2.**
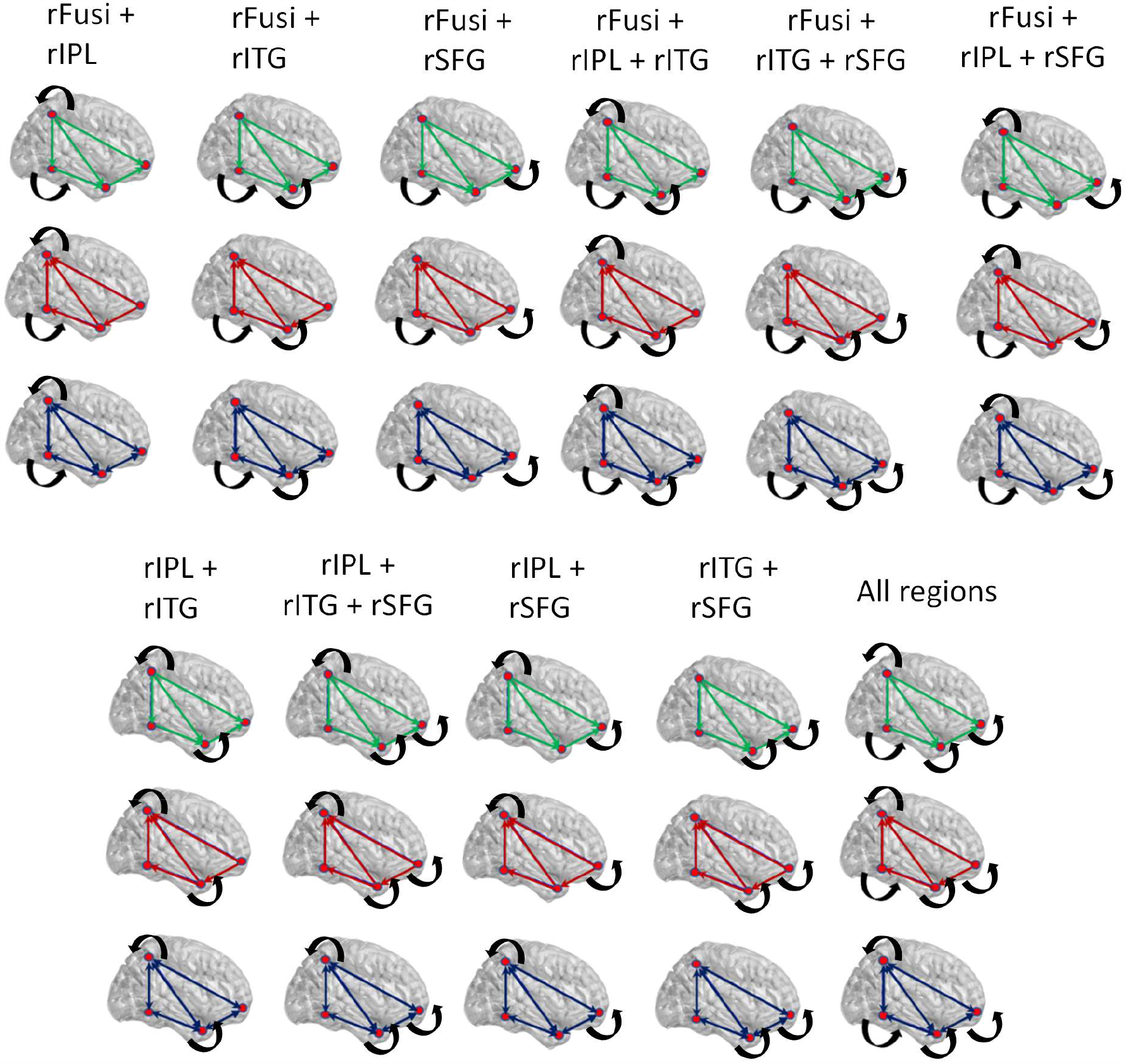
The additional 33 models within the DCM model space tested, extending on the base 15 models in Figure 3.

## Notes

### Competing Interest Statement

The authors have declared no competing interest.

